# Gene loss and relaxed selection of plaat1 in vertebrates adapted to low-light environments

**DOI:** 10.1101/2023.12.12.571336

**Authors:** Danielle H. Drabeck, Jonathan Wiese, Erin Gilbertson, Jairo Arroyave, Dahiana Arcila, S. Elizabeth Alter, Richard Borowsky, Dean Hendrickson, Melanie Stiassny, Suzanne E. McGaugh

## Abstract

Gene loss is an important mechanism for evolution in low-light or cave environments where visual adaptations often involve a reduction or loss of eyesight. The *plaat* gene family are phospholipases essential for the degradation of organelles in the lens of the eye. They translocate to damaged organelle membranes, inducing them to rupture. This rupture is required for lens transparency and is essential for developing a functioning eye. *Plaat3* is thought to be responsible for this role in mammals, while *plaat1* is thought to be responsible in other vertebrates. We used a macroevolutionary approach and comparative genomics to examine the origin, loss, synteny, and selection of *plaat1* across bony fishes and tetrapods. We show that *plaat1* (likely ancestral to all bony fish + tetrapods) has been lost in squamates and is significantly degraded in lineages of low-visual acuity and blind mammals and fish. Our findings suggest that *plaat1* is important for visual acuity across bony vertebrates, and that its loss through relaxed selection and pseudogenization may have played a role in the repeated evolution of visual systems in low-light-environments. Our study sheds light on the importance of gene-loss in trait evolution and provides insights into the mechanisms underlying visual acuity in low-light environments.

## 1. Introduction

Though evolution via sequence divergence has been a major focus of studies of adaptation, gene loss has been demonstrated to be an important mechanism of adaptive evolution, shaping both constructive and regressive adaptations [1–3]. Gene loss can occur through various mechanisms such as genetic drift and/or positive selection and is a common feature of evolution in low-light or cave environments where visual adaptations often involve a reduction or loss of eyesight [4–6]. Generally, the degradation and loss of eye-associated genes are thought to be adaptive via the loss of energy-intensive tissue or through closure of a disease-vulnerable mucus membrane in fossorial species [7–10].

Though the eye is one of the most well-studied organs, the physiological and genetic basis of many vision-related diseases remain unknown [4,11–13]. Previous comparative genomic work on cave-dwelling and fossorial lineages with eye degradation has provided vital insights into the genetic basis of eye-related gene function and disease [13–15]. In addition, insights from genomic studies of gene loss have been critical to informing convergent adaptive function [16–19].

*Plaat* family genes, which are phospholipases involved in N-acylethanolamine (NAE) biosynthesis, play a crucial role in lens clarification, as well as tumor suppression, satiation signaling, and viral translocation [20,21]. Recently, *Plaat1* was found to play a critical role in the metabolism of lens organelles in zebrafish (affecting eye clarity), as well as cardiolipin synthesis and remodeling, which may be a key novel pathway for diabetes [22]. Specifically, the protein PLAAT1 is recruited to damaged organelle membranes in the lens during development, causing the rupture and clearance of lens organelles which is required for the development of a clear lens [21]. *Plaat1* is one of 5 paralogs (*plaat1*-*5*) present in humans, and *plaat3* is thought to be the dominant phospholipase for lens organelle degradation in mice [21]. While publicly available databases (e.g. Ensembl, NCBI orthology and gene tree tools) suggest that duplication events are also apparent in primates, rodents, and shrews, the evolutionary history and function of these orthologs across mammals has not been explored. Expression and knockdown data suggest that *plaat1* is critical for eye clarity in zebrafish. Both zebrafish *plaat1* knockouts and mice *plaat3* knockouts develop cataracts, and previous work suggests that *plaat1* is conserved across vertebrates while *plaat3* is unique to mammals [20–21]. It remains unclear what role *plaat1* may play in mammalian eye development or if this gene is central to eye development in fish exclusively.

While previous evidence suggested a role for the *plaat* gene family in the evolution of visual acuity, their potential loss or pseudogenization and correlation with vision loss has not been investigated. Here we utilize a comparative phylogenetic approach to examine *plaat1* across more than 300 vertebrates and *plaat3* in mammals, including low visual acuity and blind (LVAB) mammals as well as several convergent lineages of LVAB fishes (Figure 1). Our data support the newly discovered function of *plaat1* as a vital eye-function gene showing strong evidence for relaxed selection and gene degradation in several independent lineages of LVAB fishes and mammals. These data suggest *plaat1* is important for the evolution of visual acuity in both fish and mammals and demonstrates the value of a comparative evolutionary framework to inform gene function [21,23]. While the precise impact of the *plaat* gene family on vertebrate macroevolution remains to be fully elucidated, our data show that gene loss, gene degradation, and relaxed selection of these genes is strongly associated with reduced vision in specific vertebrate clades.

**Figure 1.**
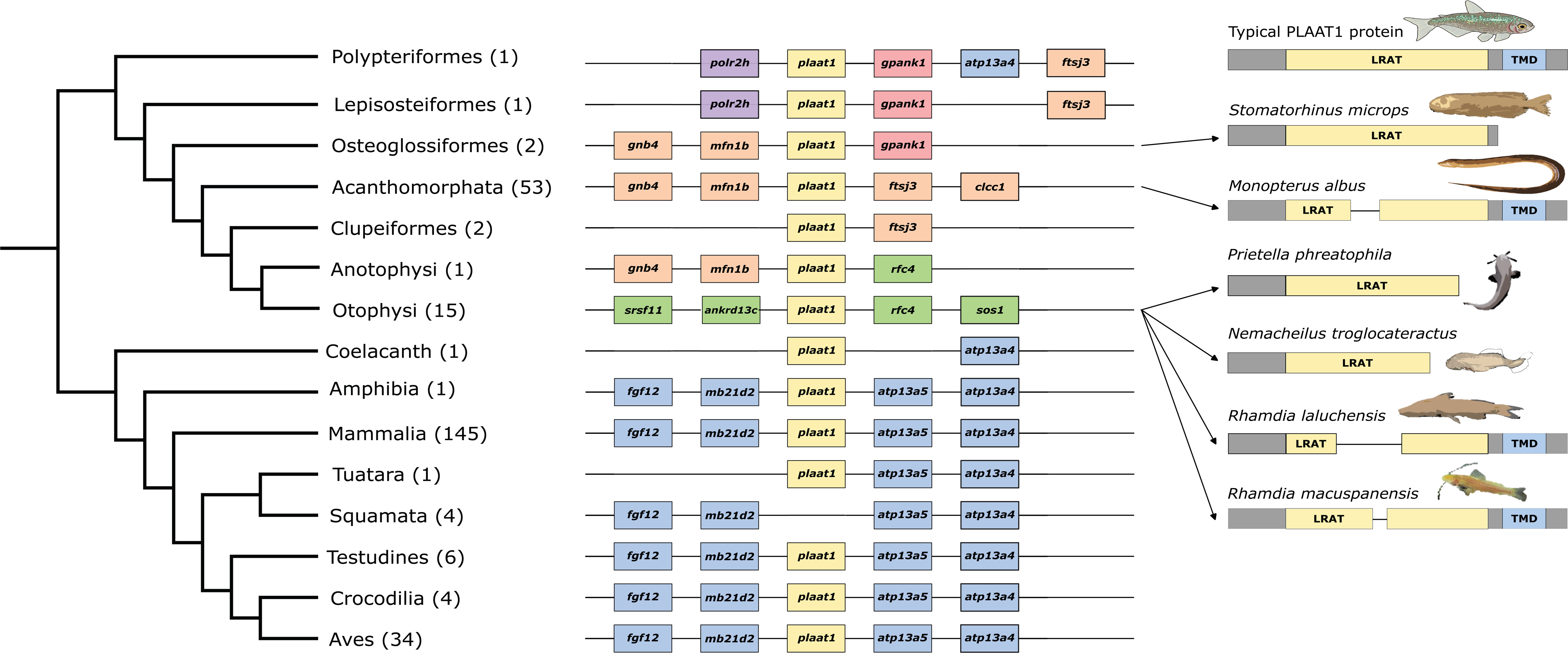
Topology of all species included in plaat1 analysis. Blue lineages indicate cave, blind, and low-visual acuity species designated as foreground for selection analyses. This dataset represents 311 species in total.

## 2. Methods

### (a) Sequence Retrieval and Alignment

In total, 311 sequences from 310 species (two *Astyanax mexicanus,* one surface and one cave) were aligned and used for downstream analysis. First, we used the NCBI ortholog database (accessed Aug 2022, https://www.ncbi.nlm.nih.gov/genome/annotation_euk/process/#references), to download one orthologous sequence of *plaat1* from each species from groups of interest—mammals, bony fishes, and sauropsids (filtering processes for orthology databases can be found here: https://www.ncbi.nlm.nih.gov/genome/annotation_euk/process/) (Supplementary file 3). Nucleotide sequences were translated into protein sequence and aligned with Clustal Omega v1.2.2 [24]. Protein alignments and DNA sequences were then used as input for PAL2NAL v14 [25] to generate protein-informed codon alignments. Second, sequences annotated as *plaat1* were pulled from 13 newly annotated cave and cryptophthalmic interstitial lithophilic river fish genomes (Supplementary Table 1), part of a concurrent project (Drabeck et al. *in prep*). To be sure that the full and correct sequence for *plaat1* was recovered, genomes were searched using BLASTN as well as an hmm-based exon capture pipeline (https://github.com/lilychughes/FishLifeExonHarvesting). Resulting hits were mapped to their respective scaffolds in GeneiousPrime 2022.2.1 (https://www.geneious.com). Scaffolds were also searched for syntenic genes using the map to reference function. Exons for *plaat1* were extracted and aligned to the RefSeq transcript from *A. mexicanus,* and accessioned into Genbank (submission ID: 2721929:OR260033-45) (AstMex3_surface GeneID: 103036287).

Sequences were filtered for quality and paralogy through a multi-step process as follows. Using unrestricted tests of selection (aBSREL, described below), we examined sequences which were found to be under positive selection via a likelihood ratio test (n=8). Since these sequences were highly divergent from the bulk of the alignment, we further scrutinized them. First, alignments were scrutinized by eye and misplaced gaps were corrected (n=3: *Xiphophorus maculatus, X. couchianus,* and *Poecilia reticulata*). We also used Ensembl’s ortholog database (release 109 - Feb 2023, https://useast.ensembl.org/info/genome/compara/homology_method.html) to check that NCBI sequences matched and were the longest available transcript, and if they were not, sequences were replaced with the longest available transcript (Ensembl n=2 *Struthio camelus, Crocodylus porosus*). If a species was not available on Ensembl, we used *plaat1* sequence from a closely related species to BLASTN against all available NCBI genomes for the species with a questionable sequence, and mapped hits as well as syntenic genes to their respective scaffolds in Geneious v8.1. Sequences for which a better sequence match was identified on the same scaffold as genes syntenic with *plaat1*, were deemed mis-annotated and removed (n=3 *Trachemys scripta elegans*, *Haliaeetus albicilla*, *Pterocles gutteralis*).

Programs used to test for selection cannot handle stop codons, thus sequences were edited to remove stop codons and contain only in-frame sequence in GeneiousPrime 2022.2.1. Three species (*Stomatorhinus microps*, *Nemacheilus troglocateractus*, *Prietella phreatophila*) had premature termination codons (PTCs) in the second or third exon after which the remnant of the gene was still discernible. To test whether signal for relaxed selection was amplified when including both the coding region and the gene remnants, we used both (selection analyses for alignments with and without gene remnants) for downstream analyses. Alignments with and without gene remnants used for analyses are available on Dryad (DOI: 10.5061/dryad.qfttdz0p0). To verify that premature stop codons were not a result of assembly errors, we performed a BLASTn on the raw reads using the entire sequence with gene remnants as a query, and included an alignment of all 500 top hits with the query in our Dryad database. TimeTree (http://www.timetree.org) was used to generate a topology for all available sequences and additional species were added in a text editor using topologies from the FishLife project (www.FishTree.org) [26]. The same process was repeated for *plaat3* (for mammals only) (Figure 3; Supplementary Data).

#### (b) Evolutionary hypothesis testing

Several nested hypotheses about the evolution of *plaat1* were tested. To account for the diversity of sequences included in this dataset, tests of selection were run using both the entire dataset as well as two subsets: 1) Actinopterygii and 2) Mammalia. Foreground lineages included LVAB mammals, cavefishes,cryptophthalmic interstitial lithophilic river fish (*Stomatorhinus microps*, *Mastacembelus brichardi*) and the Asian swamp eel (*Monopterus albus*, which has reduced eyes but is also not a cavefish) (Figure 1, Supplementary File 2). Hereafter we use simply ‘LVAB’ to reference the fish and mammal foreground species. Because plaat1 is missing entirely in squamates, and several sequences within sauropsida appear to be pseudogenized, we test whether *plaat1* was evolving under relaxed or positive selection in all sauropsids. To do this we designated sauropsids as the foreground taxa using the entire 311 sequence dataset. To test for positive selection on foreground branches, we employed an unrestricted branch site test, aBSREL as well as a gene-wide test of selection, BUSTED, in HyPhy v2.5.33 [28,29]. To test for relaxed selection, we employ the RELAX model test, specifying foreground lineages as LVAB mammals, LVAB fishes, or sauropsids (Supplementary File 2)[30]. To identify sites under positive selection in foreground lineages, we used the mixed effects model of evolution (MEME) [28,31]. All statistical tests for selection were employed within the HyPhy software suite v 2.5.33 [31].

The PLAAT proteins contain a transmembrane domain (TMD) which is vital for membrane identification and attachment, as well as a lecithin retinol acyltransferase domain (LRAT) which is responsible primarily for catalysis [20–21]. To test for enrichment of mutations that are likely to cause a loss of function, we counted for presence of a deletion or premature stop codons located in the TMD, LRAT domain, and the N-terminal (NTERM) region for both fish and mammals of *plaat1*. We used Fisher’s exact tests in R (version 4.2.3) to compare the occurrence of deletions and premature stop codons in either LVAB mammals or LVAB fishes with sighted mammals and fish, respectively.

#### (c) Confirming plaat1 gene loss in squamates

In both NCBI and Ensembl orthology databases, *plaat1* is present in mammals, archosaurs and turtles, tuataras, amphibians, lungfish, coelacanths, and cartilaginous fishes, but is missing in squamates, and therefore likely present in the most recent common ancestor (MRCA) of Gnathostomata.

Loss of *plaat1* in squamates was confirmed utilizing both automated and manual checks. First, to create a BLAST query for *plaat1*, we used BioMART’s orthology tool to compile nucleotide and amino acid sequences from five mammals spanning broad phylogenetic range (Tasmanian devil, platypus, opossum, cow, human), as well as gar and frog. This query was used to perform both BLASTN and tBLASTN searches against 90 sauropsid reference genomes (all amniotes including mammals and avian/nonavian reptiles), transcriptomes, and raw reads (Supplementary File 1). Losses were confirmed manually utilizing both NCBI and Ensembl ortholog databases (accessed Aug 2022). Finally, manual examination of syntenic regions was used to confirm gene loss and, if possible, identify gene remnants. To evaluate syntenic regions, we identified conserved genes that were directly upstream and downstream to *plaat1* using the NCBI orthology database in sauropsids (Figure 2). We then used these flanking genes to search available genomes within squamates to identify the syntenic region and recover potential gene remnants [27]. If flanking genes were clearly present but *plaat1* was missing, intergenic regions were extracted and aligned using ClustalOmega in Geneious v8 to the best quality/ closest related genome available that contained *plaat1* to recover any possible gene remnants [24].

**Figure 2.**
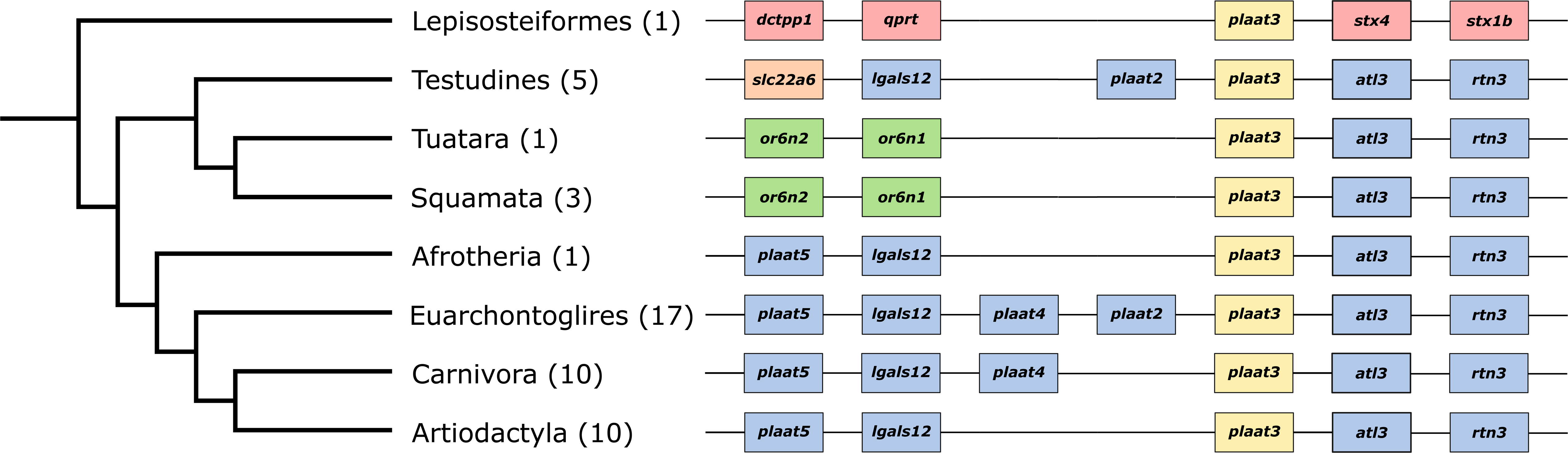
Generalized synteny of the plaat1 gene across bony fish and tetrapod species. Synteny was identified using the NCBI genome browser. Groups of species sharing the same syntenic pattern were collapsed on the phylogeny and are denoted using the largest taxonomic group of which all species are included. The number in parentheses indicates the number of species included in our analysis that fell in each group and that had a genome assembly on NCBI. Each species in each group shared the same synteny pattern, with the following exceptions. **Fish:** Osteoglossiformes: Old calabar mormyrid (Paramormyrops kingsleyae) missing mfn1b; Acanthomorphata: Ocellaris clownfish (Amphiprion ocellaris) missing clcc1 and ftsj3, Zig-zag eel (Mastacembelus armatus) missing mfn1b, Crimson soldierfish (Myripristis murdjan) missing clcc1 and ftsj3, Black rockcod (Notothenia coriiceps) missing clcc1; Otophysi: Common carp (Cyprinus carpio) missing rfc4. **Mammals:** African elephant (Loxodonta africana) missing atp13a4, Greater mouse-eared bat (Myotis myotis) missing atp13a5, Jamaican fruit bat (Artibeus jamaicensis) missing fgf12 and mb21d2, Baiji (Lipotes vexillifer) missing atp13a5, Blue whale (Balaenoptera musculus) missing atp13a5, Northern elephant seal (Mirounga angustirostris) missing fgf12. **Aves**: Helmeted guineafowl (Numida meleagris) missing fgf12 and mb21d2. PLAAT1 protein illustrations are generic and not to scale. LRAT = lecithin retinol acyltransferase domain, TMD = transmembrane domain.

## 3. Results

In brief, our comparative genomic approach recovered important losses and relaxed selection in synteny-confirmed orthologs of *plaat1*. Losses were found predominantly in blind fishes, or fishes with low visual acuity. Enrichment analysis revealed that deletions and premature stop codons were significantly enriched in functional domains for both LVAB fishes and LVAB mammals. This suggests that the *plaat1* gene has played an important role in the evolution of vision (and its subsequent loss) in many vertebrate species.

### (a) Little evidence of positive selection in lineages of LVAB fishes for *plaat1*

Neither gene-wide (BUSTED) nor branch-site (aBSREL) revealed signatures of significant positive selection across 85 teleost fish in a hypothesis-free test (Table 1). When LVAB were designated as foreground species, positive selection was detected on the branch leading to the Asian swamp eel (Table 1; Supplementary Table 4), and tests for relaxed selection with all LVAB fishes designated as foreground are marginally non-significant (*p* = 0.06). Mixed effects model of evolution (MEME) revealed no sites (p < 0.05) evolving under positive selection when LVAB fish lineages are placed in the foreground.

**Table 1.** Summary of results for various tests for diversifying and relaxed selection on plaat1 using low acuity and blind species (LVAB) mammals, LVAB fishes, and/or sauropsids as foreground taxa.LVAB fish sequences were run with and without gene remnants (GR) which included the sequence downstream of a premature stop codon which otherwise would have been within a coding exon. The designation ‘none’ means that no branch was specified as the foreground, and all branches were run and tested individually. P-values reported are corrected for multiple tests in HyPhy.

|  | Foreground Specified | aBSREL<br>(episodic positive, branch-site) | BUSTED<br>(episodic positive, gene-wide) | RELAX<br>(relaxed, gene-wide) | MEME<br>(episodic positive, site) |
| --- | --- | --- | --- | --- | --- |
| Mammals | LVAB | $p > 0.05$ | $p = 0.0208$ | $p = 0.0003$ | $p < 0.05$<br>(4 sites) |
| | none | $p > 0.05$ | $p = 0.5000$ | | |
| Fishes | LVAB | $p < 0.04^5$ | $p = 0.4777$ | $p = 0.0608^*$ | $p < 0.05$<br>(0 sites) |
| | LVAB (GR) | $p < 0.0004^4$ | $p = 0.0011$ | $p < 0.0001$ | |
| | none | $p > 0.05$ | $p = 0.4798$ | | |
| Vertebrates | LVAB | $p < 0.002^2$ | $p < 0.0001$ | | $p < 0.05$<br>(6 sites) |
| | Sauropsids | $p < 0.04^3$ | $p = 0.4441$ | $p < 0.0001$ | |
| | none | $p < 0.03^1$ | $p < 0.0001$ | | |

<sup>1</sup>*Crocodylus porosus*, p-value <0.000011, *Terrapene carolina*, p-value <0.000011, *Struthio camelus*, p-value <0.000011 *Elephantulus edwardii*, p-value = <0.000011, *Myotis davidii*, p-value = 0.02573
<sup>2</sup>*Myotis davidii*, p-value = 0.00145
<sup>3</sup>*Terrapene carolina*, p-value = <0.000010, *Crocodylus porosus*, p-value = <0.000010, *Struthio camelus*, p-value = <0.000010, *Nestor notabilis*, p-value = 0.03369
LVAB: foreground set to include all blind/low visual acuity species in that group
<sup>4</sup>*Stomatorhinus microps*, p-value < 0.000011, *Monopterus albus*, p-value = 0.0268
<sup>5</sup>*Monopterus albus*, p-value = 0.03031
\*marginal non-significance

### (b) *plaat1* exon loss and relaxed selection in LVAB fishes

*Plaat1* is a phospholipase which functions via enzymatic activity through binding to a substrate with a hyper-conserved C-terminal transmembrane region, and it has been previously established that removing this region renders PLAAT1’s phospholipase activity nonfunctional by making it unable to translocate to organelle membranes [21]. Because tests of selection are blind to insertions, deletions, and other structural changes that significantly impact gene function, we examine the instance of deletions and premature stop codons in LVAB species compared to sighted species. Among 16 fish lineages designated as foreground species based on eye reduction, one cryptophthalmic interstitial lithophilic river fish *Stomatorhinus microps*) and two cave-dwelling species (*Prietella phreatophila*, *Nemacheilus troglocataractus*) are missing the C-terminal end of *plaat1* due to premature stop codons in exon 3 (*S. microps, P. phreatophila*) or 2 (*N. troglocataractus*) (Figure 2). Across 85 teleost fish, these are the only sequences missing any portion of the C-terminal sequence (Figure 2). Two additional cave-dwelling species (*Rhamdia laluchensis* and *R. macuspanensis*) as well as the Asian swamp eel (*Monopterus albus*, which has reduced eyes but is not a cavefish) have large deletions in the middle exon (53-85aa, 67-72aa, 59-68aa, respectively) which are also unique relative to all other teleost fishes surveyed. Fisher’s exact tests confirm that deletions and premature stop codons are significantly enriched for LVAB and blind species in both the LRAT (p = 0.0001) and TMD (p = 0.0068) domains (Supplementary Figure 1).

For three fish species with significant exon loss (*S. microps, P. phreophila, N. troglocateractus*), gene remnants 3’ to a premature stop codon were identifiable by eye. The majority of teleost fishes have a PLAAT1 protein which is 166-170 amino acids in length, whereas *S. microps, P. phreophila,* and *N. troglocateractus* have lengths 136, 124, and 133 respectively, all of which are restored to a full length (166-170aa) when gene remnants are included. To test whether these regions were under relaxed selection, we employed the same test of selection with these regions included. When the sequence 3’ to premature stop codons (gene remnants) were included in the same analyses, the branch leading to the elephant fish (*S. microps*) was also identified as evolving under positive selection (aBSREL) (Table 1; Supplementary Table 5). Similarly, when gene remnants were included in gene-wide tests of positive selection (BUSTED) and relaxed selection (RELAX) signal for both were significant (p = 0.0011, p<0.0001 respectively), suggesting that regions 3’ to the premature termination codon may still be under negative selection.

### (c) *Plaat1* in low-visual acuity and blind mammals

Using the low-visual acuity and blind lineages of mammals highlighted in Figure 1, we used an array of selection tests to identify whether evolutionary trajectories of eye loss were associated with sequence-level signatures of relaxed or positive selection on *plaat1* in mammals. Though *plaat3* is thought to be most strongly involved in lens clarification in mammals, it remains unknown how important *plaat1* is to this function. Tests of branch-site selection (aBSREL) of *plaat1* did not recover evidence of positive selection for any branches when specifying low visual acuity or blind mammals as foreground species (Figure 1; Table 1). Tests for gene-wide episodic positive selection (BUSTED) were significant when identifying LVAB mammals as a foreground lineage (p=0.0208), but not significant when a foreground is not specified (p=0.5). Tests of relaxed selection using the same LVAB foreground species were also significant (Table 1; Supplementary Table 6). Similar to fishes, Mixed Effects Model of Evolution (MEME) reveals codon diversification as clustered largely at the C-terminal end (Table 1; Supplementary Table 7). Particularly notable are deletions in the TMD for the Cape golden mole and the blind mole-rat.

### (d) Plaat3

Because lens clarification via organelle degradation is thought to be more strongly influenced by *plaat3* in mammals, we would expect *plaat3* to exhibit signatures of selection (either relaxed or intensified) among LVAB mammals. To test this expectation, we aligned *plaat3* from 66 available mammals and employed tests for both positive and relaxed selection. We found largely conserved synteny across placental mammals (Figure 3) and recovered no evidence for positive selection or relaxed selection across this gene. However, both *Myotis spp.* (mouse-eared bats) and *Chrysochloris asiatica* (Cape golden mole) have very long branch lengths, and notable sequence divergence in the C-terminal region (Supplementary Figure 2).

**Figure 3.**
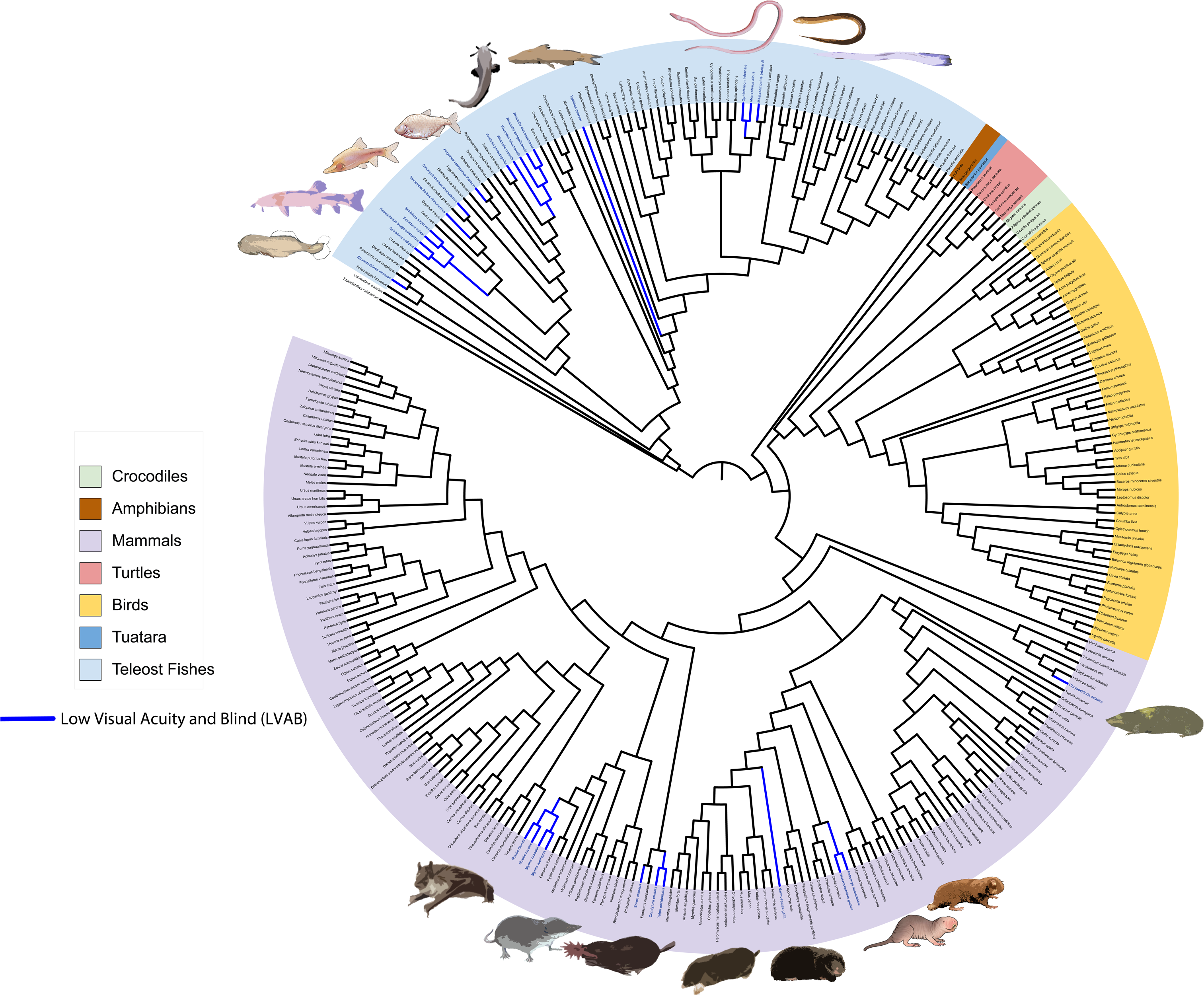
Generalized synteny of the plaat3 gene across selected vertebrate lineages. Plaat3 orthologs were identified using the Ensembl gene tree (ENSG00000176485). Synteny was identified using the NCBI genome browser. Groups of species sharing the same syntenic pattern were collapsed on the phylogeny and denoted using the largest taxonomic group of which all species are included. The number in parentheses indicates the number of species included in our analysis. Each species in each group shared the same synteny pattern, with the following exceptions: **Testudines**: Painted turtle (Chrysemys picta) missing lgals12, Three-toed box turtle (Terrapene carolina triunguis) missing plaat2; Squamata: Eastern brown snake (Pseudonaja textilis) missing or6n1; Euarchontoglires: mouse (Mus musculus) missing plaat4 and plaat2, Coquerel’s sifaka (Propithecus coquereli) missing plaat5; Carnivora: harbor seal (Phoca vitulina) missing plaat4; Artiodactyla: camel (Camelus dromedarius) missing rtn3.

### (e) Clade-wide *plaat1* gene loss in squamates

Searches of genome assemblies and raw reads, including exhaustive searches of the intergenic regions of syntenic genes suggest that *plaat1* is lost in squamates (Figure 2). Because *plaat1* is present in the gar, coelacanth, cartilaginous fish, toad, and mammals, the most parsimonious scenario for the evolution of *plaat1* is that it was present in the MRCA of gnathostomes and subsequently lost in the lineage leading to squamates.

### (f) Relaxed Selection of *plaat1* supported in Sauropsids

Branch-site tests of positive selection (aBSREL) with sauropsids specified as the foreground, which includes the stem as well as all branches within Sauropsida (Figure 1, Supplementary Figure 3), recovered several branches within sauropsids under positive selection (Table 1, Supplementary Table 2). To test for evidence of relaxed selection, we employed the HyPhy RELAX test. Specifically, RELAX compares a model where both negatively selected sites *and* positively selected sites are shifting *towards* neutrality in foreground lineages with respect to background lineages, versus a simpler model for positive selection where positively selected sites are shifting *away from neutrality*. Tests of relaxed selection using all sauropsids as foreground recovered evidence of gene-wide relaxation of selection (Supplementary Table 3), while a gene-wide test of positive selection with sauropsids placed as foreground (BUSTED) was not significant (*p*=0.44).

## 4. Discussion

Understanding the evolutionary history of gene families can shed light on the processes that drive evolutionary change such as gene duplication, gene loss, and functional divergence. By tracing the evolutionary trajectories of gene families across different lineages, we can gain insights into the evolution and loss of important biological traits over time. Though functional work is needed to better understand the importance of *plaat1* in eye development and function, our work found enrichment for deletions, premature stop codons, and relaxed selection in lineages with species with reduced eyes, as well as an ancient loss along the basal lineage of squamates. Overall, our study suggests that *plaat1* may play a vital role in evolutionary shifts to sightlessness.

### (a) *Plaat1* gene degradation in multiple species of LVAB fishes

Colonization of cave environments and the evolution of troglomorphism has occurred in over 200 species of teleost fish [46–49]. Our dataset included sixteen species with reduced eyes that represent eight independent lineages of cave colonization, two cryptophthalmic interstitial lithophilic river fish (*S. microps, M. brichardi*), and one non-cave dwelling species with greatly reduced eyes (*M. albus*) (Figure 1) [50–51]. *Plaat1* zebrafish knockouts of the entire *plaat1* gene as well as knockouts of only the C-terminal transmembrane domain (TMD), reliably produce fish with cataracts [21]. Two cavefish lineages (the blind cave loach, *N. troglocateractus*; and the Mexican blindcat, *P. phreatophila*) and the cryptophthalmic interstitial lithophilic river fish African elephant fish (*S. microps*) have premature stop codons 5’ to the TMD, and therefore likely lack a functional *plaat1* which is key for lens clarification. However, because the remainder of this gene (including the catalytic triad) remains intact, it is possible that phospholipid metabolism is conserved in some capacity (potentially avoiding negative pleiotropy). Three additional large deletions occur in three additional blind/eye-reduced fishes (the La Lucha blind catfish, *R. laluchensis*; the Olmec blind catfish, *R. macuspanensis*; and the Asian swamp eel, *M. albus*). These deletions occur in the lecithin retinol acyltransferase (LRAT) domain which plays a vital role in the availability and storage of retinol and are therefore very likely to cause large functional disruptions [52].

Gene-wide and branch-site tests of selection which did not specify a foreground across teleosts are not significant (Table 1). When LVAB fishes are specified as foreground species in branch-site tests (aBSREL), the lineage leading to *M. albus* is identified as evolving under positive selection, and when gene remnants are included in the same analysis, both *M. albus* and *S. microps* are identified as evolving under positive selection (Supplementary Table 4-5). As both species have large deletions of major functional regions (See Supplementary data annotation alignment), are identified on branches evolving under positive selection, with marginally non-significant signal for relaxed selection, we can surmise that positive selection may be contributing to loss of function. This contrasts with the case in mammals where there is a clear negative result for all branch tests of positive selection, and clearer difference in level support for relaxed (p=0.0003) vs positive selection (0.0208). T hese results suggest that selection might contribute to loss of function mutations in these species, a phenomenon that has been demonstrated for other cavefish [40].

While tests of relaxed selection specifying the same foreground are marginally non-significant (p = 0.06), they become highly significant when gene remnants are included (p < 0.0001). Together this suggests that both relaxed selection and positive selection may be important in the process of *plaat1* gene degradation and are likely acting on different sites of the same gene [40–41].

### (b) Selection for *plaat1* but not *plaat3* is relaxed in low visual acuity and blind mammals

Recovery of significant evidence for relaxed or positive selection in *plaat1* in mammals is found exclusively in tests that designate low visual acuity and blind mammals as foreground species, suggesting that *plaat1*, likely retains an important role in mammalian eye function and development. Notably, the blind Cape golden mole, *Chrysochloris asiatica*, has two coding mutations which disrupt the ultra-conserved H-box domain, thought to be vital for hydrolysis [37].

Though *plaat3* has been shown to have a vital role in lens clarification in mice, we did not see a signal of positive or relaxed selection in LVAB mammals in our dataset for *plaat3*. However, branch lengths for *plaat3* for the Cape golden mole and the little brown bat are noticeably long (though not significantly so), with sequence divergence concentrated in the C-terminal end of the protein across the TMD, which is vital for membrane translocation and lens clarification (Supplementary Figure 1 & 2).

Mammalian *plaat1* results are significant for both gene-wide episodic diversifying selection (BUSTED) as well as relaxed selection (RELAX). Though fit across models cannot be directly compared, p-values are much lower for relaxed selection, suggesting this model may be a better fit for these data. Additionally, BUSTED results identified one site with an evidence ratio >10, suggesting that a small number of positively selected sites (one) are driving the signal for positive selection [30]. Both selection and Fisher’s exact tests support the hypothesis that mammalian *plaat1* is degrading via relaxed selection in species with reduced visual acuity and blindness. These data support recent work by Partha et al. [15] which identifies HRASLS (PLA/AT) as a gene exhibiting convergent rate evolution in fossorial mammals [45].

While signal for relaxed selection and gene degradation across convergent lineages of LVAB species suggests that *plaat1* retains some functional importance in eye development in mammals, it is also possible that *plaat1* is associated with pleiotropic physiologies that are also altered in dark environments. For example, *plaat1* is known to be involved in insulin signaling pathways, cardiolipin metabolism, as well as tumor suppression, which are also known to be important in shifts to darkness [22,38,39]. Additionally*, plaat1* plays a role in p53-mediated apoptosis involved in starvation and disease, both of which are greatly modified in low-light environments [23].

### (c) Loss of *plaat1* in squamates

*Plaat1* is present in the tuatara (sister group of squamates) and absent in both lizards and snakes, suggesting that it was lost along the branch leading to squamates. Shifts to fossorial niches and subsequent reduction of eye-related genes occurred at several points across both modern and ancient snakes and lizards [35]. Recent molecular work has shown that loss of eye related genes is apparent across the squamate lineage, including in the squamate ancestor [35]. The apparent loss of *plaat1* uncovered here agrees with recent work and identifies a novel pathway of eye reduction in ancient squamates [35].

### (d) Relaxed selection of *plaat1* in sauropsids

Because *plaat1* is lost in squamates, and several sauropsids appear to have highly degraded *plaat1* sequences, we investigated whether relaxed selection was pervasive in all squamates, indicating a more widespread loss of function, redundancy, or alternate function in this larger group. Results of selection tests indicate significant relaxed selection across sauropsids in *plaat1* (Table 1, Supplementary Table 3). Sequence divergence across sauropsids is quite variable, with likely non-functional variants and potentially pseudogenized *plaat1* convergently arising in the tuatara, common box turtle, saltwater crocodile, kea, and the red-throated loon (Supplementary Figure 3) [36]. Further investigation should focus on the role of *plaat1* in sauropsids, and whether this apparent divergence is driven by convergent physiological functional adaptation, ancient loss of function, or by neofunctionalization of sauropsid-specific paralogs which may have rendered *plaat1* redundant.

### (e) Other important physiologies

While we focus here on the newly discovered lens clarification function that is apparent for *plaat1*, it is important to note that *plaat1* has several pleiotropic physiological interactions, which we do not fully understand [20–22,36,39,52]. *Plaat* family proteins are important for tumor suppression, NAPE biosynthesis (hormones released by the small intestine into the bloodstream to process fat), obesity, and peroxisome formation [20–22,36,39,52,53]. Particularly noteworthy is *plaat1*’s role in facilitating viral translocation, making viral coevolution another strong potential driver of convergent gene loss and/or degradation in vertebrates [53]. When *plaat1* is suppressed in mouse and human cell lines, NAPE levels decrease, suggesting that *plaat1* is an important N-acetyltransferase [20]. *Plaat1* deletions in humans are associated with Polland Syndrome, a disorder which causes missing or underdeveloped muscles on one side of the body [52,54].

*Plaat3*’s absence or divergence outside of mammals is also noteworthy, as it plays vital roles in obesity, cancer invasion, and vitamin A storage [55–57]. Remarkably, in the course of this work, we observed the venomous mammal *Sorex araneus* had more than 30 duplications of *plaat3*, which is also present in the salivary transcriptome of another venomous shrew, *Blarina brevicaudata,* which suggests that *plaat3* might have been recruited into the venom proteome in shrews.

## 5. Conclusions

There is a lack of research on the impact of the *plaat* gene family on vertebrate macroevolution. Specifically, our work suggests that this gene family may have played a crucial role in the evolution of visual acuity. We found a significant association between *plaat1* increased sequence divergence (due to relaxed or positive selection), premature stop codons, and deletions in key functional domains across 12 lineages of low visual acuity and blind mammals, 16 lineages of low-light adapted fishes, and a complete loss of *plaat1* in squamates. We identified several cases of convergent relaxed selection within sauropsids that have notably reduced visual acuity or are night vision specialists (e.g., kiwi and saltwater crocodiles, respectively). However, convergent relaxed selection in other groups (e.g. ostrich), with high visual acuity, suggest that the functional significance of *plaat1* within sauropsids remains unknown, and requires further exploration. Though there are at least 5 paralogs known in humans, and public databases suggest duplication events elsewhere in the mammalian tree, further investigation of ortholog and paralog function are needed to shed light on how labile lens clarification function is among copies. A better understanding of how these enzymes have diversified, and how different species have coped with losses and gains of function may be crucial to understanding disease pathways (e.g., eye disease, insulin signaling, tumor suppression, and viral infection) and developing strategies for human disease intervention[14].

## Acknowledgments

This work was supported by the National Institute of Health Teaching Research Educators in Minnesota grant (grant #K12GM119955), and the University of Minnesota Undergraduate Research Opportunities Program. Additional financial support for this research was provided to JA by the Universidad Nacional Autónoma de México (UNAM) through a “Programa de Apoyo a Proyectos de Investigación e Innovación Tecnológica” (PAPIIT) grants (IA200517 and IA202119) and by the Consejo Nacional de Ciencia y Tecnología (CONACyT) through a “Ciencia Básica” grant (A1-S-28293), and to SEA and MLJS by National Science Foundation award DEB-1655227. The authors would like to extend their gratitude to Rachel Moran, Matt Holding, Matt Winn and Sergei Pond for their feedback and suggestions and help with data visualization and analyses, as well as the Minnesota Supercomputing Institute (MSI) for use of computing resources and support. Special thanks to Emma Roback for feedback, suggestions, and the use of her original digital art. The authors would like to thank Dean Hendrickson and the San Antonio Zoo for aiding in acquiring the tissues necessary for this and accompanying genomic work.

## Data Accessibility

The data generated in this work including sequence alignments, species topologies, and scaffolds are stored in the Dryad repository which can be accessed with this link DOI: 10.5061/dryad.qfttdz0p0

## Supplementary Figure Legends

**Supplementary Figure 1.**
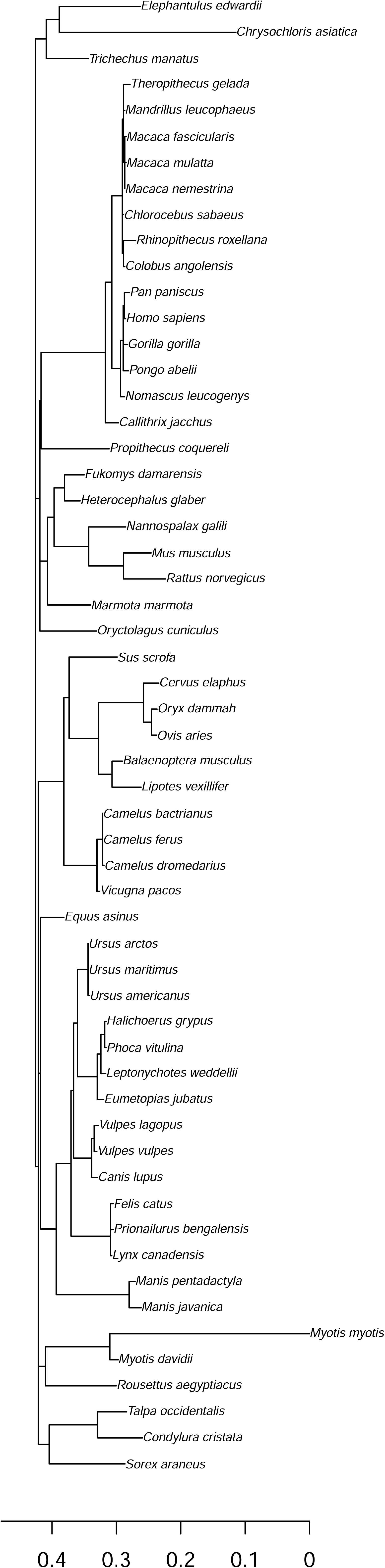
Bar plot depicting the number of deletions or premature stop codons (PSCs) in each region of *plaat1* for (A) sighted versus low visual acuity and blind (LVAB) mammals, and (B) sighted versus LVAB fishes. Genes are counted only once if they have one or more PSC or deletion to reduce bias caused by degraded genes having multiple deletions/disruptions. Numbers under bars indicate total number of sequences (species) that fell into that category. P-values derived from Fisher’s exact tests.

**Supplementary Figure 2.**
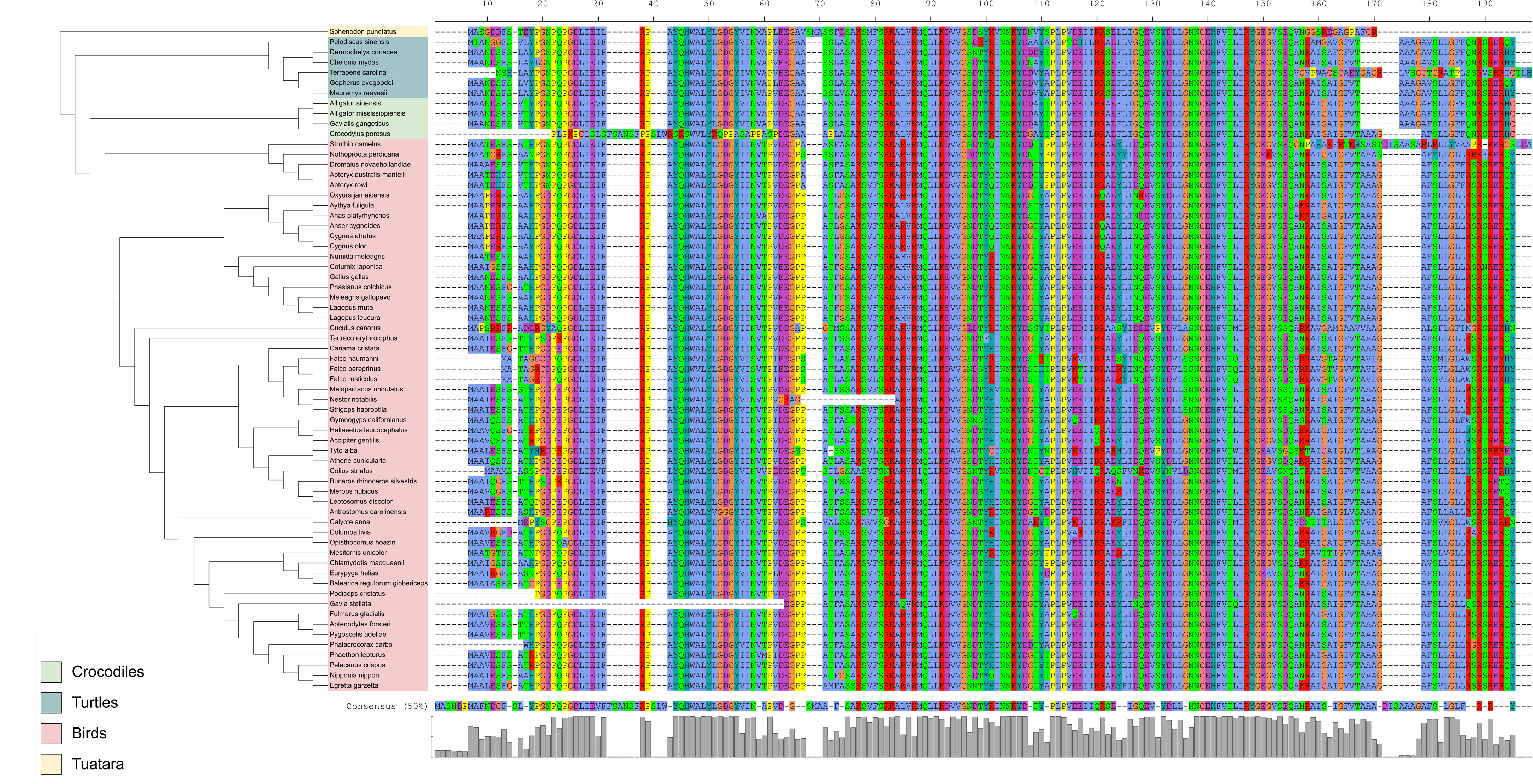
Gene tree of *plaat3* for all mammals included in selection analyses, generated using aBSREL Full adaptive model (Smith et al. 2015).

**Supplementary Figure 3.**
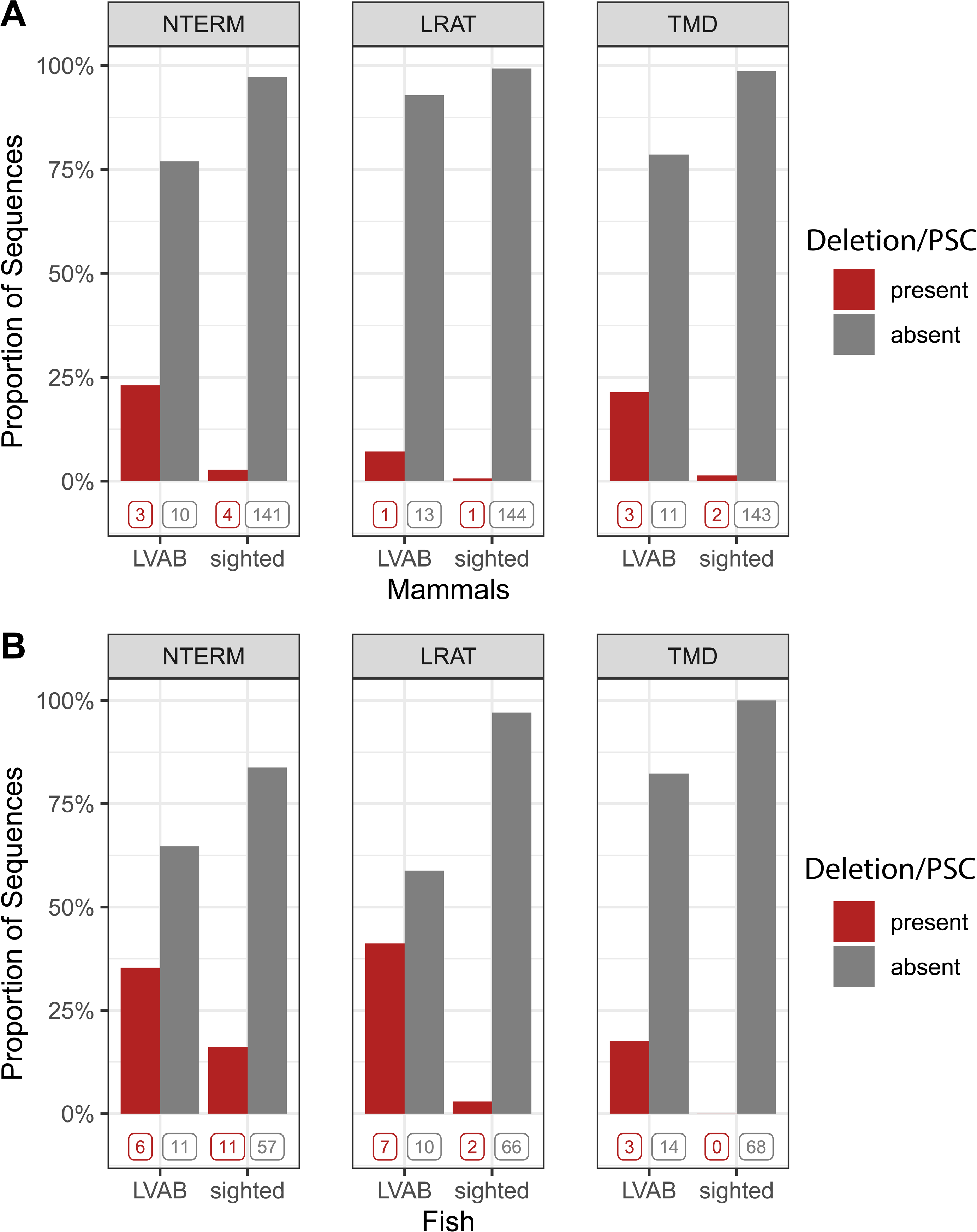
Topology of all sauropsids used in analyses next to the protein alignment of *plaat1* for each species. Image generated in iTOL.

