## Supplementary Table for "Gene loss and relaxed selection of plaat1 in vertebrates adapted to low-light environments"

**Supplementary Table 2.** aBSREL analysis of *plaat1* with sauropsids in the foreground. aBSREL found evidence of episodic diversifying selection on 4 out of 554 branches. A total of 117 branches were formally tested for diversifying selection. Significance was assessed using the Likelihood Ratio Test at a threshold of p ≤ 0.05, after correcting for multiple testing. Significance and number of rate categories inferred at each branch, and percent of sites inferred per rate category are provided in the detailed results table. Branches with signatures of selection <0.05 are included.

| **Name** | **B** | **LRT** $\chi2$ | **Test p-value** | **Uncorrected p-value** | **ω distribution over sites** |
| --- | --- | --- | --- | --- | --- |
| *Crocodylus_porosus*  *(SaltWater Crocodile)* | <0.00001 | 95.6038 | <0.00001 | <0.00001 | ω_1_ = 0.501 (83%) ω_2_ = 224 (17%) |
| *Struthio_camelus*  *(Ostrich)* | <0.00001 | 53.8968 | <0.00001 | <0.00001 | ω_1_ = 0.0209 (80%) ω_2_ = 9090 (20%) |
| *Terrapene_carolina*  *(Box Turtle)* | <0.00001 | 97.4557 | <0.00001 | <0.00001 | ω_1_ = 0.0215 (72%) ω_2_ = 3850 (28%) |
| *Nestor_notabilis*  *(Kiwi)* | <0.00001 | 14.0957 | 0.0337 | 0.0003 | ω_1_ = 0.204 (98%) ω_2_ = 612 (1.9%) |

**Supplementary Table 3.** RELAX model fits using all 311 vertebrate *plaat1* sequences with Sauropsids in the foreground. Test for selection relaxation (K = 0.05) was significant (p <0.0001, LR = 75.23).

| **Model** | ***log* L** | **# params** | **AIC_c_** | **Branch set** | **ω_1_** | **ω_2_** | **ω_3_** |
| --- | --- | --- | --- | --- | --- | --- | --- |
| General descriptive | -29255.5 | 1127 | 60793.4 | Shared | 0.00 (9.48%) | 0.00 (86.00%) | 22.11 (4.52%) |
| RELAX alternative | -29846.5 | 574 | 60848.3 | Reference | N/A | 0.36 (24.96%) | 3.54 (2.86%) |
|  |  |  |  | Test | N/A | 0.95 (24.96%) | 1.06 (2.86%) |
| RELAX null | -29884.1 | 573 | 60921.5 | Reference | 0.01 (70.51%) | 0.38 (25.63%) | 2.76 (3.86%) |
|  |  |  |  | Test | 0.01 (70.51%) | 0.38 (25.63%) | 2.76 (3.86%) |
| RELAX partitioned descriptive | -29846.4 | 578 | 60856.2 | Reference | 0.01 (73.78%) | 0.31 (22.73%) | 3.19 (3.49%) |
|  |  |  |  | Test | 0.00 (73.05%) | 0.82 (3.94%) | 1.02 (23.01%) |

**Supplementary Table 4.** aBSREL results from plaat1 using 85 teleost fish with cavefish identified as the foreground. Branches with signatures of selection <0.05 are included.

| **Branch** | **B** | **LRT** | **Test p-value** | **Uncorrected p-value** | **ω distribution over sites** |
| --- | --- | --- | --- | --- | --- |
| *Monopterus albus* | <<0.000011 | 10.7535 | 0.0303 | 0.0016 | ω_1_ = 0.236 (96%) ω_2_ = 3850 (3.7%) |

**Supplementary Table 5.** aBSREL results from plaat1 using 85 teleost fish with cavefish identified as the foreground, including portions of the gene 3’ to a premature stop codon. Branches with signatures of selection <0.05 are included.

| **Branch** | **B** | **LRT** | **Test p-value** | **Uncorrected p-value** | **ω distribution over sites** |
| --- | --- | --- | --- | --- | --- |
| *Stomatorhinus_microps* | *<<0.000011* | *81.9893* | *<<0.000011* | *<<0.000011* | *ω1 = 0.0111 (82%)*  *ω2 = 100000 (18%)* |
| *Monopterus_albus* | *<<0.000011* | *10.8895* | *0.0268* | *0.0015* | *ω1 = 0.235 (96%)*  *ω2 = 3800 (3.8%)* |

**Supplementary Table 6.** Model fits for RELAX using 135 mammals identifying reduced acuity/blind mammals as foreground branches. Test for selection relaxation (K = 0.10) was significant (p = 0.000, LR = 14.45).

| **Model** | ***log* L** | **#. params** | **AIC_c_** | **Branch set** | **ω_1_** | **ω_2_** | **ω_3_** |
| --- | --- | --- | --- | --- | --- | --- | --- |
| General descriptive | -10767.9 | 571 | 22701.6 | Shared | 0.00 (69.89%) | 0.00 (30.01%) | 981.97 (0.10%) |
| RELAX alternative | -10906.6 | 296 | 22411.6 | Reference | 0.00 (62.96%) | 0.68 (37.04%) | N/A |
|  |  |  |  | Test | 0.14 (62.96%) | 0.96 (37.04%) | N/A |
| RELAX null | -10913.8 | 295 | 22424.0 | Reference | 0.01 (61.45%) | 0.67 (38.55%) | N/A |
|  |  |  |  | Test | 0.01 (61.45%) | 0.67 (38.55%) | N/A |
| RELAX partitioned descriptive | -10903.8 | 300 | 22414.2 | Reference | 0.01 (64.18%) | 0.68 (35.82%) | N/A |
|  |  |  |  | Test | 0.39 (76.36%) | 0.39 (23.30%) | 42.40 (0.34%) |

**Supplementary Table 7:** MEME on all mammals episodic positive/diversifying selection at 4 sites with p-value threshold of 0.05.

| **Site** | **Partition** | **LRT** | **p-value** | **# branches under selection** | **Total branch length** | **MEME LogL** | **FEL LogL** |
| --- | --- | --- | --- | --- | --- | --- | --- |
| **36** | 1 | 9.49 | 0.00 | 1.00 | 0.00 | -38.20 | -34.73 |
| **171** | 1 | 9.15 | 0.00 | 0.00 | 0.00 | -76.43 | -76.43 |
| **152** | 1 | 6.37 | 0.02 | 0.00 | 0.00 | -75.67 | -73.01 |
| **140** | 1 | 4.50 | 0.05 | 1.00 | 0.00 | -38.50 | -37.33 |
